# Programmable RNA Targeting using CasRx in Flies

**DOI:** 10.1101/2020.04.03.023606

**Authors:** Anna Buchman, Dan J. Brogan, Ruichen Sun, Ting Yang, Patrick Hsu, Omar S. Akbari

## Abstract

CRISPR-Cas genome editing technologies have revolutionized the fields of functional genetics and genome engineering, but with the recent discovery and optimization of RNA-targeting Cas ribonucleases, we may soon see a similar revolution in the study of RNA function and transcriptome engineering. However, to date, successful proof-of-principle for Cas ribonuclease RNA targeting in eukaryotic systems has been limited. Only recently has successful modification of RNA expression by a Cas ribonuclease been demonstrated in animal embryos. This previous work, however, did not evaluate endogenous expression of Cas ribonucleases and only focused on Cas ribonuclease function in early developmental stages. A more comprehensive evaluation of this technology is needed to assess its potential impact in the field. Here we report on our efforts to develop a programmable platform for RNA-targeting using a Cas ribonuclease, CasRx, in the model organism *Drosophila melanogaster*. By genetically encoding CasRx in flies, we demonstrate moderate transcript targeting of known phenotypic genes in addition to unexpected toxicity and lethality. We also report on the off-target effects following on-target transcript cleavage by CasRx. Taken together, our results present the current state and limitations of a genetically encoded programmable RNA-targeting Cas system in *Drosophila melanogaster*, paving the way for future optimization of the system.

## Introduction

The development of CRISPR as a programmable genome engineering tool has revolutionized the life sciences by providing transformative applications for both medicine and biotechnology [1]. While much of the recent focus has been on exploiting CRISPR technologies to target DNA, recent findings that certain CRISPR systems can also be programmed to target RNA have suggested new possibilities for CRISPR technologies in transcriptome engineering [2–4]. For example, one recent advancement was the engineering and biochemical characterization of CasRx as a compact single-effector Cas enzyme that exclusively targets RNA with superior efficiency and specificity as compared to RNA interference (RNAi) [4]. In human cells, CasRx demonstrated highly efficient on-target gene reduction with limited off-target activity, making it a potential tool for gene reduction. However, this technology has yet to be comprehensively adapted for facile use in other systems (although see [5]), such as *Drosophila melanogaster* (flies), which are a common tool for exploring new biological questions and developing novel bioengineering tools *in vivo.* Non-RNAi based techniques for reducing gene expression (without permanently altering the genome) in animals would provide for a more flexible technique to modulate gene expression in a biologically relevant way.

CasRx belongs to the Cas13 family of RNA-targeting Cas enzymes, a group of highly specific ribonucleases [4,6]. Though these enzymes possess promiscuous RNase activity resulting in cleavage of non-target RNA [2,4,7–9], a possible drawback for applying Cas13 ribonuclease-based transcriptome engineering technologies, they may serve as a starting point for optimizing these RNA-targeting platforms for *in vivo* applications. For example, RNA-targeting CRISPR technologies could enable the development of robust gene silencing techniques in animals in which RNAi poorly functions [4,10]. Another potential application may involve using RNA-targeting CRISPR technologies to engineer mosquito populations resistant to infection with RNA viruses. Numerous RNA viruses of global medical importance, such as dengue, Zika, and chikungunya virus, are transmitted by one species of *Aedes* mosquito. Engineering this mosquito population with virus resistance may be a tool to reduce disease transmission [11]; however, no current technologies have successfully targeted all of these viruses simultaneously [12–16]. RNA-targeting CRISPR systems may provide a platform to reduce the spread of mosquito-borne viruses by targeting viral RNA genomes in a programmable manner. Therefore, it is of high priority to further understand the utility of RNA targeting CRISPR systems in relevant model organisms.

RNAi-based approaches are the current standard for transcriptome modification in *Drosophila.* This technology has increased our understanding of the function and regulation of many genes [10,17–20]. Despite these efforts, limitations of RNAi technologies are numerous and consequently advancements in this field have stagnated in recent years. High false negative rates, particularly in highly expressed genes due to insufficient small RNA expression [10,17,21], high false positive rates due to positional or off-target effects [22–25] and the difficulty of simultaneously targeting multiple genes, has limited the research capabilities of RNAi. Coexpression of Dicer2 can reduce false negative rates, but also may increase the prevalence of false positives [10,17] and further complicates the method. Ideally, an RNA targeting system should be easily programmable, not require expression of multiple factors, and should work in a simplified manner. CasRx, like other CRISPR systems, is easily programmable [26,27] and is capable of targeting nearly any coding gene, but unlike other Cas13 enzymes, it lacks a protospacer flanking sequence requirement [4] making it more versatile for programmable targeting. Additionally, CasRx is a simplified RNA-targeting system as it requires no additional helper enzymes to efficiently target and degrade RNA [4]. For these reasons, the CasRx ribonuclease is a practical starting point for establishing a single-effector RNA-targeting platform for *in vivo* gene reduction studies. Here we report the first use of a CasRx-mediated RNA-targeting system in flies. We show that separately encoding CasRx and guide RNA arrays (gRNA^array^) in the genome promotes robust expression of these elements throughout development. Furthermore, we demonstrate that binary genetic crosses with ubiquitous and tissue-specific CasRx- and gRNA^array^-expressing fly lines can produce clear, highly penetrant phenotypes and by using RNA sequencing (RNAseq) we demonstrate that CasRx is capable of moderate targeted transcript reduction at various stages of fly development, albeit with various degrees of off-target activity. Moreover, we also found that CasRx expression and targeting was often toxic and resulted in unexpected lethality indicating further optimization will be necessary for this to be a versatile tool for *Drosophila* genetics.

## Results

### Genetically encoding CasRx in flies

To genetically determine the RNA-targeting capabilities of CasRx, *in vivo*, we engineered flies encoding two versions of the CasRx ribonuclease, the active enzyme and a catalytically inactive negative control (dCasRx). We did this by generating transgenes that use a broadly expressing ubiquitin (Ubiq) promoter [28] to drive expression of either CasRx (Ubiq-CasRx) or dCasRx (Ubiq-dCasRx) (Fig. S1). CasRx exhibits two distinct RNase activities for processing its cognate gRNA^array^ and cleaving target RNA. Because we wanted our negative control to still bind target RNA and efficiently process the gRNA^array^, we eliminated programmable RNA cleavage in dCasRx by inactivating the positively charged catalytic residues of the HEPN motifs [4]. We established these transgenic lines by integrating each transgene site-specifically using an available *ϕ*C31 docking site located on the 2nd chromosome (attp40w) (Fig. S1, Table S1). These flies were viable, though we were unable to generate homozygotes for either CasRx or dCasRx, presumably due to high levels of ubiquitous ribonuclease expression. While homozygotes are desirable because, when outcrossed, all progeny would receive a copy of the transgene, we were still able to assess CasRx activity by maintaining these stocks as heterozygotes balanced over the chromosome Curly-of-Oster (CyO), which ensures a non-lethal expression level of CasRx while retaining the transgene (Table S1). To genetically measure the efficacy of programmable RNA targeting, we targeted three genes known to produce visible phenotypes when expression is disrupted, including: *white* (*w*), *Notch (N),* and *yellow (y)* [29–32]. To target these genes with CasRx, we designed a gRNA^array^ for each gene driven by a ubiquitously expressed polymerase-3 U6 (U6:3) promoter [33,34] (Fig. S1, Table S1). Each array consisted of four transcript-targeting spacers (30 nt in length) each positioned between CasRx-specific direct repeats (36 nt in length) with a conserved 5’-AAAAC motif designed to be processed by either CasRx or dCasRx [4] (Fig. S1). For each gRNA^array^, we site-specifically integrated the transgene at an available *ϕ*C31 docking site located on the 3rd chromosome (site 8622) and established a homozygous transgenic line (Fig. S1, Table S1).

### Programmable RNA targeting of endogenous target genes

To assess the efficacy of programmable RNA targeting by CasRx, we conducted bidirectional genetic crosses between homozygous gRNA^array^ (+/+; gRNA^array^/gRNA^array^) expressing flies crossed to either Ubiq-CasRx (Ubiq-CasRx/CyO; +/+) or Ubiq-dCasRx (Ubiq-dCasRx/CyO; +/+) expressing flies (Fig. 1A). When crossed to Ubiq-CasRx, we were able to obtain highly-penetrant (100%) expected visible eye pigmentation disruption phenotypes exclusively in transheterozygotes (Ubiq-CasRx/+; gRNA^array^ /+) for *w* suggesting that CasRx exhibits programmable on-target RNA cleavage capabilities (Fig. 1B,C, Table S2). However, while we expected Mendelian transheterozygote inheritance rates to be 50%, the recorded inheritance rates were significantly lower than expected (12.9%), suggesting some degree of unexpected toxicity leading to lethality (Fig. 1B, Fig. S2, Table S2). Moreover, when targeting *y* or *N,* Ubiq-CasRx transheterozygotes (Ubiq-CasRx/+; gRNA^array^ /+) were 100% lethal and did not develop beyond the second instar larval stage (Fig. S3 A,C). This was expected for *N* as there are many examples of lethal alleles for this gene [35–37], however mutations in *y* should be recessive viable with defective chitin pigmentation producing yellow cuticle phenotypes [38]. Furthermore, we were unable to obtain phenotypes in transheterozygotes (Ubiq-dCasRx/+; gRNA^array^ /+) from our negative control crosses using all arrays tested, indicating that a catalytically active form of the ribonuclease is necessary for phenotypes to be observed (Fig. 1C). Taken together, these results indicate that the catalytically active form of the CasRx ribonuclease can generate phenotypes, although unexpected toxicity which resulted in lethality (only in the presence of the CasRx and the array) were also observed.

**Fig. 1.**
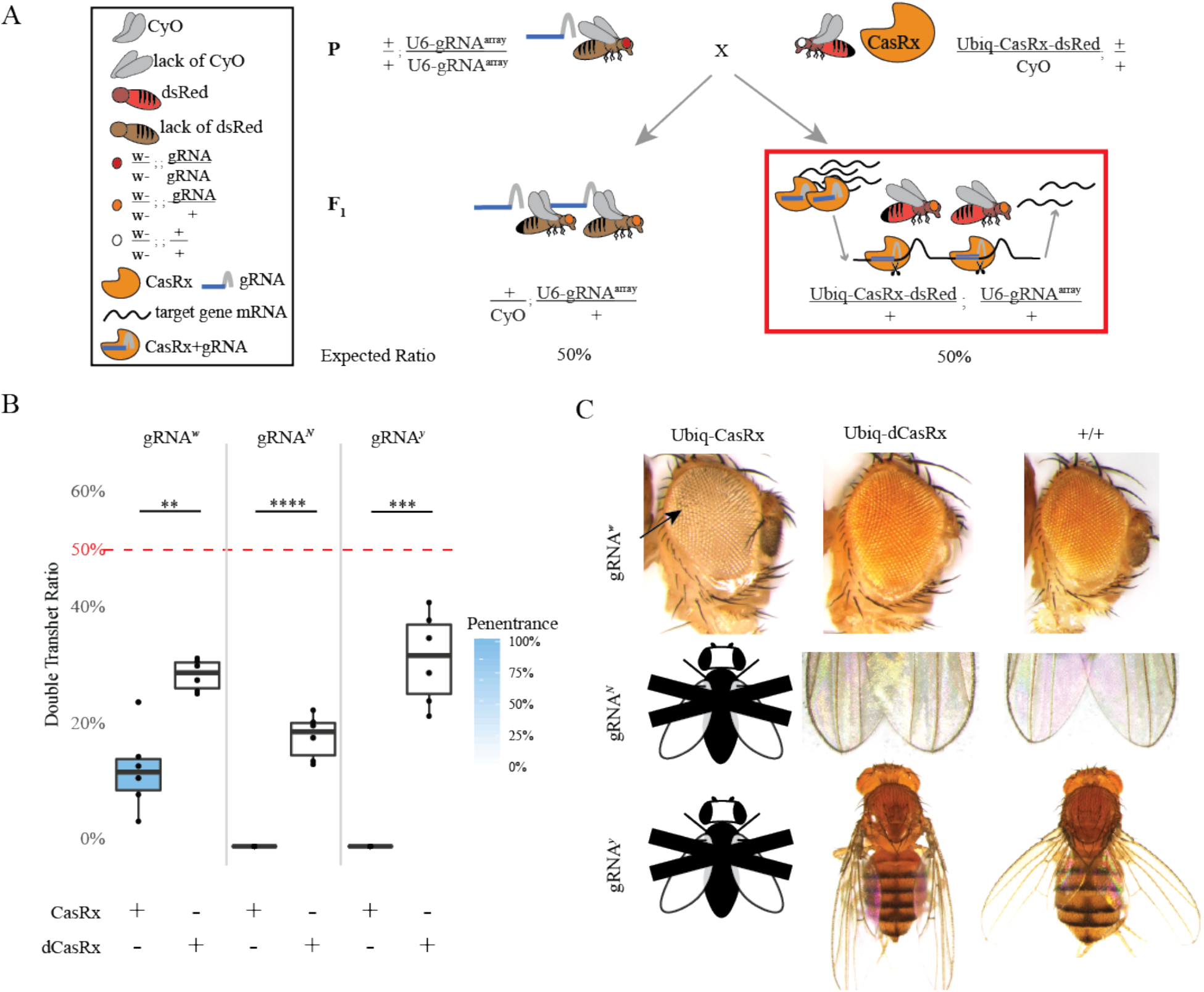
Genetic assessment of programmable CasRx-mediated transcript reduction in flies. (A) Representative genetic crossing schematic to generate transheterozygotes. (B) Inheritance and penetrance rates of transheterozygous flies inheriting both Ubiq-CasRx (or Ubiq-dCasRx) and gRNA^array^ corresponding to the red box in panel A. Phenotype penetrance rate is depicted by blue shading in the box plot. Significant differences in inheritance between CasRx and dCasRx groups were observed in all 3 groups (p values: gRNA^*w*^ = 0.00135; gRNA^*N*^ = 0.00006; gRNA^*y*^ = 0.00016). (C) Brightfield images of transheterozygous flies with representative phenotypes for each cross. Corresponding genotype for each image is dictated by the combination of constructs on top of the columns and the side of the rows. Arrows point to tissue necrosis in the eye. Black and white fly with “X” represents lethality phenotype where no transheterozygote adults emerged.

### Tissue-specific RNA targeting by CasRx

Given the above toxicity when ubiquitously expressed, we next explored the efficiency of CasRx when expression was restricted to specific cell types and tissues. We leveraged the classical binary Gal4/UAS system which enables targeted gene expression [39]. To develop this system, we generated two transgenes using the UASt promoter [39] to drive expression of either CasRx (UASt-CasRx) or dCasRx (UASt-dCasRx) as a negative control (Fig. S1). These transgenes were integrated site-specifically using a *ϕ*C31 docking site located on the 2nd chromosome (site 8621), and these stocks were homozygous viable (Fig. S1, Table S1). To activate CasRx expression in specific tissues, we used available Gal4 driver lines that restricted expression to either the eye (GMR-Gal4) [40] or the wing and body (yellow-Gal4) [41] (Table S1). These lines were crossed to the same homozygous gRNA^array^ lines described above targeting *w, y*, or *N* (Fig. S1, Table S1). To test this system, we performed a two-step genetic crossing scheme to generate F_2_ triple transheterozygotes (either UASt-CasRx/+; gRNA^array^/Gal4 or UASt-dCasRx/+; gRNA^array^/Gal4) (Fig. 2A). This consisted of initially crossing homozygous gRNA^array^ (gRNA^array^/gRNA^array^) expressing flies to heterozygous, double-balanced UASt-CasRx (UASt-CasRx/Cyo; TM6/+) flies, or for the negative control, heterozygous, double-balanced UASt-dCasRx (UASt-dCasRx/Cyo; TM6/+) flies. The second step was to cross the F_1_ transheterozygote males expressing both a CasRx ribonuclease and the gRNA^array^ (UASt-CasRx/+; gRNA^array^/TM6 or UASt-dCasRx/+; gRNA^array^/TM6) to respective homozygous Gal4 driver lines to generate F_2_ triple transheterozygotes (UASt-CasRx/+; gRNA^array^/Gal4 or UASt-dCasRx/+; gRNA^array^/Gal4) to be scored for phenotypes (Fig. 2A).

**Fig. 2.**
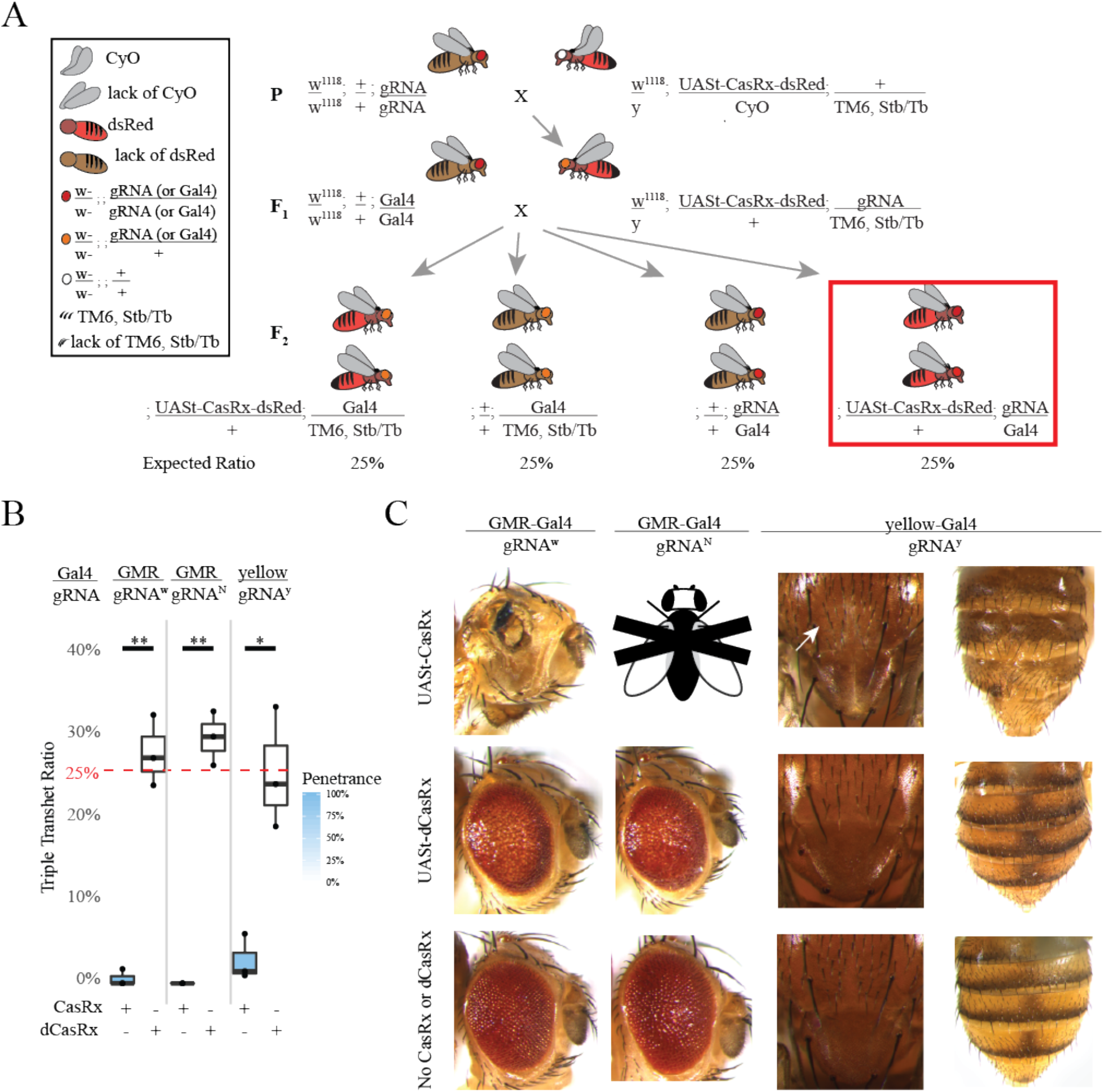
CasRx-mediated target transcript reduction in restricted tissue types using the binary Gal4/UAS system. (A) Representative genetic crossing schematic. (B) Inheritance rates of triple transheterozygous flies inheriting three transgenes (UASt-CasRx or UASt-dCasRx, gRNA^array^, and Gal4-driver), corresponding to flies highlighted in the red box in panel A. Significant differences in inheritance between CasRx and dCasRx groups were observed in all 3 gene targets (gRNA^*w*^, p = 0.00595; gRNA^*N*^. p = 0.00402; gRNA^*y*^ p = 0.02205) (C) Phenotypes of the triple transheterozygous flies. The white arrow identifies chitin pigment reduction in the thorax resulting from *y* targeting. Black and white fly with “X” represents a lethal phenotype with no live adults able to be scored or imaged.

From these crosses, our results indicated that tissue-specific expression of CasRx can indeed result in phenotypes, though this was also accompanied by tissue-specific cell death or organismal lethality, similar to previous observations of ubiquitous CasRx expression described above. For example, of the expected 25% Mendelian inheritance rates from the F_1_ cross between gRNA^w^ (UASt-CasRx/+; gRNA^w^/TM6) and GMR-Gal4 (+/+; GMR-Gal4/GMR-Gal4), we observed survival of only 0.57% viable F_2_ triple transheterozygotes (UASt-CasRx/+; gRNA^w^/GMR-Gal4), all of which displayed severe eye specific pigmentation and morphology phenotypes (Fig. 2B,C, Fig. S4, Table S3). This gRNA^*w*^ F_2_ triple transheterozygote inheritance rate was significantly lower than the corresponding negative control F_2_ triple transheterozygote (UASt-dCasRx/+; gRNA^w^/GMR-Gal4) inheritance rate, which was closer to the expected 25% Mendelian inheritance (27.6%) (Fig. S4, Table S3). Moreover, using the same Gal4 driver (GMR-Gal4), a significant difference in inheritance was also observed for *N* targeting, which resulted in 100% lethality in F_2_ triple transheterozygotes (UASt-CasRx/+; gRNA^N^/GMR-Gal4) compared to the 29.3% inheritance rate for the negative control F_2_ triple transheterozygotes (UASt-dCasRx/+; gRNA^N^/GMR-Gal4) (Fig. 2B,C, Fig. S4, Table S3). Finally, when targeting *y* using the yellow-Gal4 driver (+/+; y-Gal4/y-Gal4), we observed marginal chitin pigment reduction as small patches of yellow cuticle on the thorax and abdomen in F_2_ triple transheterozygotes (UASt-CasRx/+; gRNA^y^/y-Gal4) (Fig. 2C, arrows) which is a phenotype that would be expected when *y* is disrupted. Similar to crosses described above, the F_2_ triple transheterozygote (UASt-CasRx/+; gRNA^y^/y-Gal4) inheritance was significantly lower (2.67%) when compared to the control F_2_ triple transheterozygote (UASt-dCasRx/+; gRNA^y^/y-Gal4) inheritance (25.2%), indicating a substantial degree of lethality (Fig. 2B,C, Fig. S4, Table S3). In all negative control crosses, phenotypes were not observed in F_2_ triple transheterozygotes (UASt-dCasRx/+; gRNA^array^/Gal4) again indicating that functional catalytic residues of the HEPN motifs are necessary for generating phenotypes observed (Figure 2B,C, Table S3). Taken together, these results demonstrate that tissue-specific expression of CasRx using the classical Gal4/UAS approach can result in phenotypes. However, as seen in the ubiquitous binary approach above, toxicity and lethality phenotypes were also observed again limiting the utility of the system.

### Criteria for CasRx-mediated phenotypes

Because CasRx on-target cleavage resulted in unexpected lethality we set out to determine the importance of target sequence availability to CasRx-mediated lethality. To do so, we opted to target a gene that is not natively expressed in flies. Therefore, we generated a GFP reporter assay to assess the necessity of a target sequence in CasRx-mediated lethality while simultaneously visualizing on-target transcript reduction. We designed a binary GFP reporter construct consisting of both a CasRx gRNA^array^ targeting *GFP* along with GFP expression driven by the broadly expressing OpIE2 promoter (gRNA^*GFP*^) (Fig 3, Fig. S1, Table S1) [42]. We established a homozygous transgenic line (+/+; gRNA^*GKP*^-OpIE2-GFP/gRNA^*GFP*^-OpIE2-GFP) by site-specifically integrating the construct at an available *ϕ*C31 docking site located on the 3rd chromosome (site 8622) (Fig. S1, Table S1). To test for GFP transcript targeting, we performed bidirectional crosses between homozygous flies expressing gRNA^*GFP*^ (+/+; gRNA^*GFP*^-OpIE2-GFP/gRNA^*GFP*^-OpIE2-GFP) to heterozygous Ubiq-CasRx expressing flies (Ubiq-CasRx/CyO; +/+) or heterozygous Ubiq-dCasRx expressing flies (Ubiq-dCasRx/CyO; +/+) as a negative control (Fig. 3A). With this assay, we observed 100% larval lethality for F_1_ transheterozygotes (Ubiq-CasRx/+; gRNA^*GFP*^-OpIE2-GFP/+), while larval lethality was completely eliminated in F_1_ transheterozygote controls (Ubiq-dCasRx/+; gRNA^*GFP*^-OpIE2-GFP/+). Lethality was also observed regardless of the maternal or paternal deposition of CasRx (Fig. 3B, Table S2). Given that GFP expression was also visible in larvae, we monitored the development of the F_1_ progeny and observed that Ubiq-CasRx transheterozygotes survived only to the first instar developmental stage, but not beyond (Fig. S3). Given this survival, we imaged first instar transheterozygote (Ubiq-CasRx/+; gRNA^*GFP*^-OpIE2-GFP/+) larvae and observed near-complete reduction in GFP expression for Ubiq-CasRx transheterozygote larvae as compared to Ubiq-dCasRx transheterozygote (Ubiq-dCasRx/+; gRNA^*GFP*^-OpIE2-GFP/+) control larvae indicating robust CasRx mediated target transcript (GFP) reduction (Fig. 3C). Taken together, these results suggest that CasRx possesses programmable RNA-targeting activity, and the lethality is dependent upon the availability of a target sequence as well as enzymatic RNA cleavage mediated by the positively charged residues of CasRx HEPN domains.

**Fig. 3.**
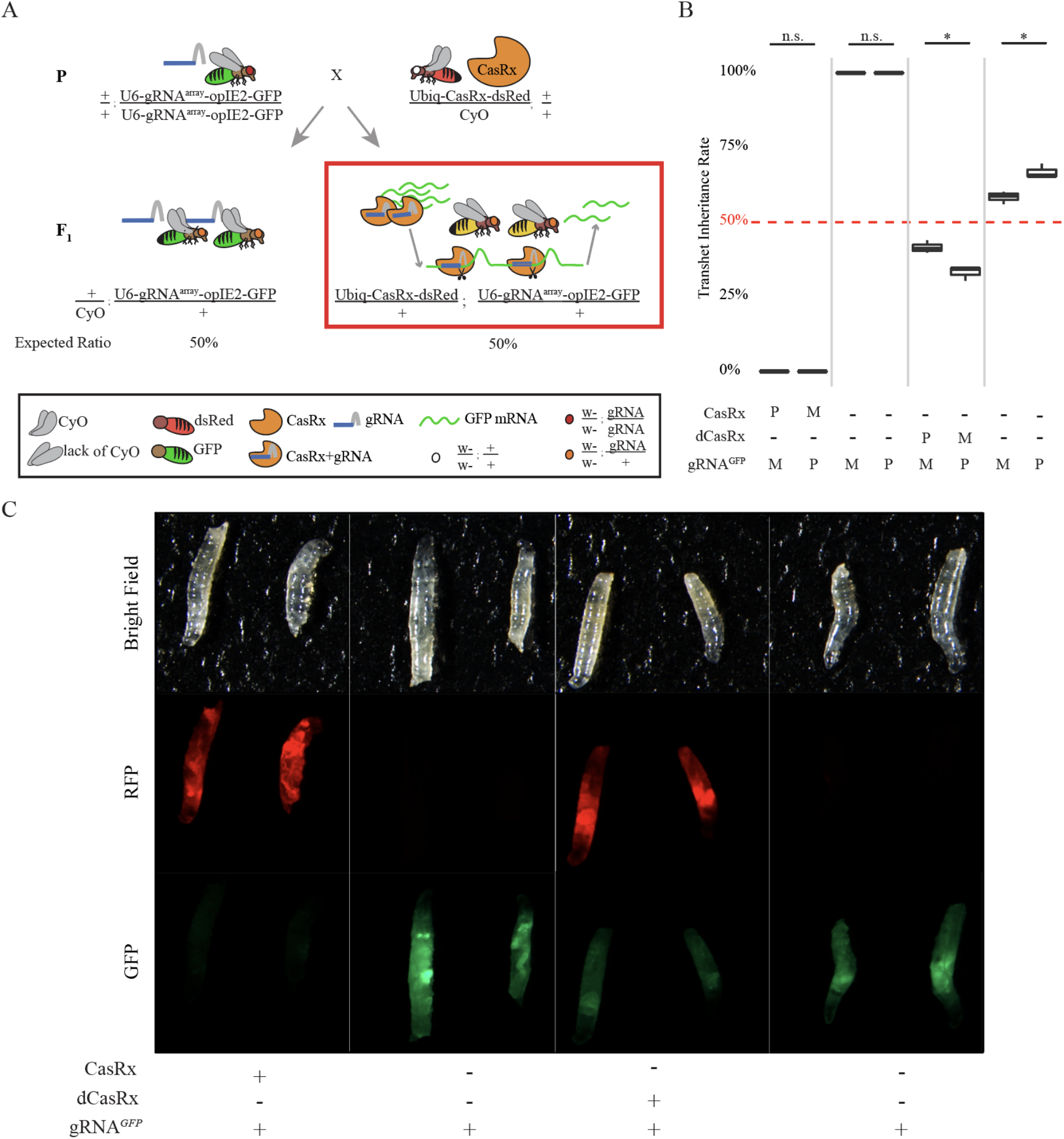
Robust CasRx-mediated reduction of GFP. (A) Representative bidirectional genetic crossing schematic. (B) Box plot of transheterozygote inheritance rates resulting from bidirectional crosses between Ubiq-CasRx (or Ubiq-dCasRx) and gRNA^*GFP*^-OpIE2-GFP flies (M = maternal inheritance of CasRx; P = paternal inheritance of CasRx). (C) Images of F_1_ larvae from paternal crosses demonstrating significant reduction in GFP expression for transheterozygous larvae expressing both Ubiq-CasRx and gRNA^*GFP*^-OpIE2-GFP compared to control transheterozygotes expressing Ubiq-dCasRx and gRNA^*GFP*^-OpIE2-GFP or without expressing a CasRx protein. (Leftright) Ubiq-CasRx/gRNA^*GFP*^ transheterozygous larvae, heterozygous gRNA^*GFP*^ larvae from Ubiq-CasRx cross, Ubiq-dCasRx/gRNA^*GFP*^ transheterozygous larvae, heterozygous gRNA^*GFP*^ larvae from Ubiq-dCasRx cross.

### Quantification of CasRx-mediated on/off-target activity

We next aimed to quantify both the on- and potential off-target transcript reduction rates. To do this, we analyzed all gRNA^array^ target genes from our binary crosses producing either highly-penetrant, visible phenotypes (*w*) or lethal phenotypes (*N*, *y*, and *GFP)* (Table S4). To do so, we implemented whole-transcriptome RNAseq analysis comparing F_1_ Ubiq-CasRx transheterozygotes (Ubiq-CasRx/+; gRNA^array^ /+) to control F_1_ Ubiq-dCasRx transheterozygotes (Ubiq-dCasRx/+; gRNA^array^ /+) (Fig. 1A red box, Fig. 3A red box, Table S4). Using the available transcriptome data of *Drosophila melanogaster* (modENCODE) [43], we extracted total RNA stages of development when high transcript expression levels were expected for each target gene with the exception of *GFP,* where we sequenced first instar larvae (Fig. S5, Table S4). In total, we analyzed 24 samples (Table S4). From our bioinformatic analysis, we found reduced target transcript expression (Fig. 4A, B). For example, of the four target genes, CasRx was able to target and significantly reduce the target transcript expression of three genes compared to dCasRx controls: *N*, *y*, and *GFP* (Fig. 4B, Table S5-S12). Although we did not observe significant transcript reduction targeting *w* we did consistently observe relative expression reduction by comparing Ubiq-CasRx samples to Ubiq-dCasRx controls, indicating some degree of on-target reduction which likely contributes to the phenotypes observed (Fig. 4B, Table S5-S12). We also quantified the number of genes with significantly misexpressed transcripts by comparing Ubiq-CasRx to Ubiq-dCasRx using DESeq2 [44] (Fig. 4A, red dots, Table S8-S12). Across all gene targets, we observed some evidence of potential off-target activity, which we define as significantly misexpressed genes between CasRx and dCasRx samples. The observed off-target activity was demonstrated by significant changes in the gene-expression levels of numerous nontarget transcripts. The number of significantly differentially expressed non-target transcripts in each group are: 253 (*w*), 300 (*N*), 41 (*y*), and 5880 (*GFP*), representing 1.4% (*w*), 1.7% (*N*), 0.23% (*y*), and 33% *(GFP)* of the total transcripts (Fig 4A, C, Table S8-S12). Taking a closer look at the gene-expression profiles of the four gene targets, we found that a total of 6,082 transcripts (out of 17,779) displayed significant expression level changes in at least one of the six CasRx-expressing groups compared to their corresponding dCasRx-expressing control group (Table S8-S12). Among the 6,082 misexpressed transcripts, 5,722 transcripts are affected by only one of the four genes targeted when CasRx is present, 334 transcripts are affected by two gene targets, 20 transcripts are affected by three gene targets, and 6 transcripts are affected by four gene targets simultaneously (Table S8-S12). This quantitative analysis of CasRx-mediated transcript reduction provides evidence of CasRx ribonuclease capabilities in flies, while also identifying potential off-target effects resulting in significantly misexpressed non-target genes.

**Fig. 4.**
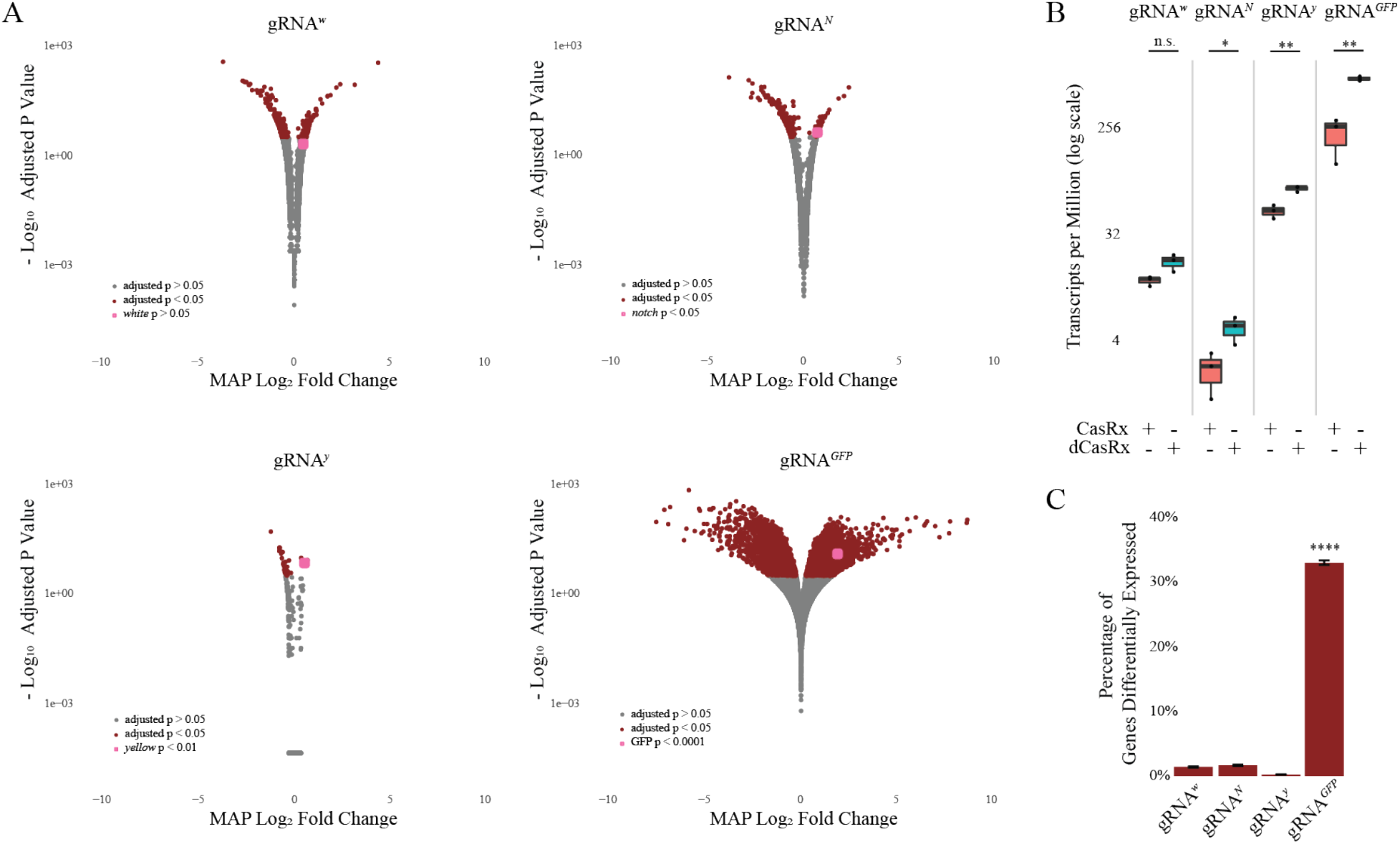
Quantification of CasRx-mediated on/off target activity. (A) Maximum *a posteriori* (MAP) estimates for the logarithmic fold change (LFC) of transcripts. DESeq2 pipeline was used for estimating shrunken MAP LFCs. Wald test with Benjamini-Hochberg correction was used for statistical inference. Grey dots represent transcripts not significantly differentially expressed between Ubiq-CasRx and Ubiq-dCasRx group (p > 0.05). Red dots represent transcripts significantly differentially expressed between CasRx and dCasRx group (p < 0.05). Pink dot identifies the respective CasRx target gene for each analysis (p value indicated in th/by inset). (B) Transcript expression levels (TPM) of transcripts targeted with CasRx or dCasRx. Student’s t-test was used to calculate significance (p values: w = 0.07, N = 0.04, y = 0.006, GFP = 0.008). (C) Percentage of transcripts significantly differentially expressed resulting from CasRx cleavage. A pairwise two-sample test for independent proportions with Benjamini-Hochberg correction was used to calculate significance.

## Discussion

Our results demonstrate that CasRx has some potential for programmable RNA targeting in flies as we did observe some expected phenotypes for each target transcript including *GFP (lethality, reduction in GFP expression)*, *N (lethality), y (lethality, yellow-patches on cuticle and thorax)* and *w (White eyes and necrosis in eyes for Gal4)*. Specifically, RNA targeting was demonstrated with ubiquitous, inducible and tissue specific CasRx expression systems against native and synthetic RNA targets, which are prerequisites for enabling comprehensive studies of gene function. However, we did also consistently observe both cellular toxicity from the ubiquitous expression of CasRx and dCasRx as we could not generate homozygous strains for either, and unexpected lethality and tissue necrosis, presumably due collateral off target effects which have been a feature previously observed for many CRISPR ribonucleases including CasRx [2,4,7–9,45]. Nevertheless, in both bidirectional and Gal4/UAS crosses, we were able to obtain visible phenotypes as well as quantitative evidence (e.g., RNAseq data demonstrating a reduction in target gene expression) indicating that the CasRx is capable of targeting and degrading target RNA in flies. It is interesting to note that, for one of the targeted genes (*w*), while the observed phenotype indicated consistent on-target transcript reduction, DESeq2 analysis did not reveal significant on-target reduction, which may be due to the timing of sample collection for RNAseq since expression levels of these genes vary over development.

Notwithstanding, we were able to obtain expected visual phenotypes in addition to significant on-target CasRx mediated transcript reduction for three of the targeted genes: *y*, *N*, and *GFP*. Interestingly, transheterozygotes (Ubiq-CasRx/+; gRNA^array^ /+) for *y*, *N*, and *GFP* also had many other misexpressed non-target genes, possibly indicating that target cleavage results in increased collateral off-target activity that is detrimental to development as these individuals were adult lethal. For example, targeting *GFP*, a non-essential gene, produced the largest number of misexpressed genes as well as the most significant fold change in expression compared to all other gene targets analyzed. Additionally, because *Gadd45,* a gene involved in cellular arrest and apoptosis in *Drosophila melanogaster [46],* was also significantly misexpressed in 4 samples (*w*, *N*, *y*, and *GFP)* (Table S8-12), it is possible that CasRx cleavage may result in an increased level of misexpressed genes leading to lethality or cellular apoptosis. Moreover, for the off target analysis for most of the target genes less than 300 (1.7%) other genes were misexpressed, however for GFP we found 5880 (33%) of genes misexpressed and it remains unclear whether this is a result of guide specific off-target or simply bystander cleavage (i.e. collateral off targeting).

Through this study, we identified two main factors contributing to CasRx-mediated lethality: (i) the catalytic activity of the CasRx HEPN domains, as lethality and tissue necrosis phenotypes were eliminated in dCasRx compared to CasRx crosses, and (ii) the presence of the target transcript resulting in on-target cleavage, as lethality was only observed when crossing Ubiq-CasRx-expressing flies to gRNA^GFP^-expressing flies. These results recapitulate previous mechanistic analyses of CasRx and other Cas13 ribonucleases, demonstrating that collateral off-target activity following targeted transcript cleavage is a native feature of Cas13 ribonucleases [2,4,7–9,45]. While this feature may not be desirable for generating tools for targeting specific transcripts of genes, this may be useful for generating sensors that get activated in response to a target transcript (e.g. viral target) that could result in activation of a marker or even organismal lethality.

Taken together, further optimization will be required to increase the CasRx on-target cleavage rates and decrease cellular toxicity and off-target effects, but this is the first demonstration of a genetically encoded programmable RNA-targeting Cas system in *Drosophila melanogaster*. In the future, optimization of the strength and timing of CasRx expression could mitigate some of the off-target associated lethality in this system. Stricter and more tunable regulation of CasRx expression may also improve phenotype penetrance as it appears to be dosage dependent in both our system and other CasRx systems [5]. For example, the phenotypes of *y* varied by their expression, with ubiquitous expression of CasRx resulting in a Ubiq-CasRx/+; gRNA^y^ /+ lethal phenotype and embryo and wing and body specific expression mitigated lethality phenotype seen in Ubiq-CasRx expression. Optimization of gRNA design may further improve these systems as CasRx gRNAs have been shown to have variable knockdown efficiency [5,47]. Nevertheless, this is an important first step towards making transcriptome engineering a viable *in vivo* technology and provides a foundation for future experiments to mitigate the off-target and toxic attributes of the enzyme to make a new, viable tool in the expanding gene-editing toolbox.

## Methods

### Design and assembly of constructs

To select the CasRx target sites, target genes were analyzed to identify 30-nucleotide (nt) regions that had no poly-U stretches greater than four base pairs, had GC base content between 30% and 70%, and were not predicted to form strong RNA hairpin structures. Care was also taken to select target sites in RNA regions that were predicted to be accessible, such as regions without predicted RNA secondary or tertiary structure (Fig. S6). All RNA folding/hairpin analysis was performed using mFold [48]. For transgenic gRNA arrays, four targets per gene were selected to ensure efficient targeting. We assembled four CasRx- and dCasRx-expressing constructs under the control of one of two promoters: Ubiquitin-63E (Ubiq) or UASt (Ubiq-CasRx, Ubiq-dCasRx, UASt-CasRx, UASt-dCasRx) using the Gibson enzymatic assembly method [49]. A base vector (Addgene plasmid #112686) containing piggyBac and an attB-docking site, the Ubiq promoter fragment, SpCas9-T2A-GFP, and the Opie2-dsRed transformation marker was used as a template to build all four constructs [33]. To assemble constructs OA-1050E (Addgene plasmid #132416, Ubiq-CasRx) and OA-1050R (Addgene plasmid #132417, Ubiq-dCasRx), the SpCas9-T2A-GFP fragment was removed from the base vector by cutting with restriction enzymes SwaI and PacI and then was replaced with CasRx and dCasRx fragments amplified with primers 1050E.C3 and 1050E.C4 (Table S13) from constructs pNLS-RfxCas13d-NLS-HA (pCasRx) and pNLS-dRfxCas13d-NLS-HA (pdCasRx) [4], respectively. To assemble constructs OA-1050L (Addgene plasmid #132418, UASt-CasRx) and OA-1050S (Addgene plasmid #132419, UASt-dCasRx), the base vector described above (Addgene plasmid #112686) was digested with restriction enzymes NotI and PacI to remove the Ubiq promoter and SpCas9-T2A-GFP fragments. Then the UASt promoter fragment and CasRx or dCasRx fragments were cloned in. The UASt promoter fragment was amplified from plasmid pJFRC81 [50], with primers 1041.C9 and 1041.C11 (Table S13). The CasRx and dCasRx fragments were amplified with primers 1050L.C1 and 1050E.C4 (Table S13) from constructs pCasRx and pdCasRx, respectively.

We designed four constructs, each carrying a four-gRNA array: OA-1050G (Addgene plasmid #132420), OA-1050I (Addgene plasmid #132421), OA-1050J (Addgene plasmid #133304), OA-1050Z4 (Addgene plasmid #132425), targeting transcripts of *white, Notch, GFP,* and *yellow,* respectively. To generate a base plasmid, OA-1043, which was used to build all constructs carrying the four-gRNA array, Addgene plasmid #112688 containing the *miniwhite* gene as a marker, an attB-docking site, a *D. melanogaster* polymerase-3 U6 (U6:3) promoter fragment, and a gRNA fragment with a target, scaffold, and terminator sequence was digested with restriction enzymes AscI and XbaI to remove the U6:3 promoter and gRNA fragments. Then, the U6:3 promoter fragment was amplified from the same Addgene plasmid #112688 with primers 1043.C1 and 1043.C23 (Table S13) and was cloned back using Gibson enzymatic assembly. To generate constructs OA-1050G, OA-1050I, and OA-1050Z4, plasmid OA-1043 was digested with restriction enzymes PstI and NotI. Then, a fragment that contained arrays of four tandem variable gRNAs (comprised of a 36-nt direct repeat [DR] and a 30-nt spacer) corresponding to different target genes followed by an extra DR and a seven-thymine terminator was synthesized and subcloned into the digested backbone using Gene Synthesis (GenScript USA Inc., Piscataway, NJ). To generate construct OA-1050J, a fragment containing arrays of four tandem variable gRNAs targeting *GFP* with an extra DR and a seven-thymine terminator followed by the OpIE2-GFP marker was synthesized and subcloned into the above digested OA-1043 backbone using Gene Synthesis (GenScript USA Inc., Piscataway, NJ). We have also made all plasmids and sequence maps available for download and/or order at Addgene (www.addgene.com) with identification numbers listed in Fig. S1 and Table S1.

### Fly genetics and imaging

Flies were maintained under standard conditions at 26°C. Embryo injections were performed at Rainbow Transgenic Flies, Inc. (http://www.rainbowgene.com). All CasRx and dCasRx-expressing lines were generated by site-specifically integrating our constructs at available *ϕ*C31 integration sites on the 2nd chromosome (sites 8621 [UAS/-(d)CasRx] and attp40w [Ubiq-(d)CasRx]). Homozygous lines were established for UASt-CasRx and UASt-dCasRx, and heterozygous balanced lines were established for Ubiq-CasRx and Ubiq-dCasRx (over Curly of Oster: CyO). All gRNA^array^-expressing lines were generated by site-specifically integrating constructs at an available *ϕ*C31 integration site on the 3rd chromosome (site 8622). Homozygous lines were established for all gRNA^array^-expressing flies.

To genetically assess the efficiency of CasRx ribonuclease activity, we bidirectionally crossed Ubiq-CasRx- and Ubiq-dCasRx-expressing lines to gRNA^array^-expressing lines at 26°C. F_1_ transheterozygotes were scored for the inheritance and penetrance of observable phenotypes. Embryo, larvae, and pupae counts were preceded by crossing male Ubiq-CasRx- or Ubiq-dCasRx-expressing flies to female gRNA^array^-expressing flies. Flies were incubated at 26°C for 48 h with yeast to induce embryo laying. Flies were then transferred to embryo collection chambers containing yeast-smeared grape-juice plates and were incubated at 26°C overnight (16 h). The grape-juice plates were then removed, the embryos were counted, and the grape-juice plates were incubated for 24 h at 26°C. Total larvae and transheterozygote larvae were then counted, and the grape-juice plates were transferred to jars and incubated at 26°C. Once all larvae reached the pupal stage, total and transhet pupae were counted. Finally, total adult flies and total adult transheterozygotes were counted 20 days post initial lay. Each genetic cross was set using 5 ♂ and 10 ♀ (paternal CasRx) or 4 ♂ and 8 ♀ (maternal CasRx) flies in triplicate.

To investigate the tissue-specific activity of CasRx, we designed a two-step crossing scheme to generate F_2_ triple transheterozygotes (Fig. 2A). First, we crossed double-balanced UASt-CasRx- or UASt-dCasRx-expressing flies (♂) to homozygous gRNA^array^-expressing flies (♀) to generate F_1_ transheterozygote males carrying the TM6-balancer chromosome. The F_1_ transheterozygote males carrying TM6 were then crossed with a Gal4–driver-expressing line. F_2_ triple transheterozygous inheritance and phenotype penetrance was scored based on visible phenotypes manifesting in flies F_2_ flies with red eyes, a lack of the TM6 balancer chromosome, and dsRed expression. Marked by the presence of dsRed (for UASt-CasRx or UASt-dCasRx), red eyes (to mark the gRNA), and the lack of TM6, F_2_ triple transheterozygotes inheritance and phenotype penetrance was scored. Each cross was set using 1 ♂ and 10 ♀ flies in triplicate. The flies were imaged on the Leica M165FC fluorescent stereomicroscope equipped with a Leica DMC4500 color camera. Image stacks of adult flies were taken in Leica Application Suite X (LAS X) and compiled in Helicon Focus 7. Stacked images were then cropped and edited in Adobe Photoshop CC 2018.

### Illumina RNA-Sequencing

The total RNA was extracted from F_1_ transheterozygous flies at different developmental stages based on the expression data available through modENCODE analysis (Fig. S5). For gRNA^*w*^ flies, transheterozygous adult heads were removed one day after emerging and were frozen at −80°C. For gRNA^*y*^ flies, the flies were incubated in vials for 48 h with yeast to induce embryo laying. The flies were then transferred to embryo collection chambers containing yeast-smeared grape-juice plates and incubated at 26°C for 3 h. The flies were then removed, and the embryos were aged on the grape-juice plates (gRNA^*y*^ = 17 h, 17–20 h total). The embryos were removed from the grape-juice plates, washed with diH_2_O, and frozen at −80°C. The gRNA^*N*^ and gRNA^GFP^ flies were incubated in vials with yeast for 48 h to induce embryo laying. The flies were then transferred to a new vial and allowed to lay overnight (16 h). The adults were removed, and the vials were incubated at 26°C for 24 h. Transheterozygote first instar larvae were then picked (based on dsRed expression) and frozen at −80°C. For all samples, the total RNA was extracted using Qiagen RNeasy Mini Kit (Qiagen 74104). Following extraction, the total RNA was treated with Invitrogen Turbo™ DNase (Invitrogen AM2238). The RNA concentration was analyzed using a Nanodrop One^*c*^ UV-vis spectrophotometer (ThermoFisher ND-ONEC-W). The RNA integrity was assessed using an RNA 6000 Pico Kit for Bioanalyzer (Agilent Technologies #5067-1513). The RNA-seq libraries were constructed using NEBNext Ultra II RNA Library Prep Kit for Illumina (NEB #E7770) following the manufacturer’s instructions [51]. The libraries were sequenced on an Illumina HiSeq2500 in single read mode with a read length of 50 nt and a sequencing depth of 20 million reads per library following the manufacturer’s instructions. Base calls were performed with RTA 1.18.64 followed by conversion to FASTQ with bcl2fastq 1.8.4. We sequenced and analyzed three replicates for all CasRx and dCasRx samples. In total, we sequenced and analyzed 24 samples: 12 CasRx experimental samples and 12 dCasRx control samples. All raw sequencing data can be accessed at NCBI SRA submission ID SUB6818910 (BioProject PRJNA600654).

### Bioinformatics - Quantification and differential expression analysis

Reads were mapped to the *Drosophila melanogaster* genome (BDGP release 6, GenBank accession GCA_000001215.4) supplemented with cDNA sequences of CasRx and GFP using the default parameters of STAR aligner [52] with the addition of the “--outFilterType BySJout” filtering option and “--sjdbGTFfile Drosophila_melanogaster.BDGP6.22.97.gtf” GTF file downloaded from ENSEMBL. The expression levels were determined with featureCounts [53] using “-t exon -g gene_id -M -O --fraction” options. The raw transcript counts were normalized to transcripts per million (TPM), which were calculated from count data using the “addTpmFpkmToFeatureCounts.pl” Perl script (Supp. File addTpmFpkmToFeatureCounts.pl). The raw count and TPM data are available in Table S5–S6. To further explore CasRx-induced differential gene expression profiles, we used the maximum *a posteriori* (MAP) method with the original shrinkage estimator in the DESeq2 pipeline to estimate the transcript logarithmic fold change (LFC) (44). The Wald test with the Benjamini-Hochberg correction was used for statistical inference. The analysis script can be found in Supp. “deseq2_analysis.R”, and the analyzed results are in Table S8–S12. Per the DESeq2 analysis requirements, some values are shown as NA for the following reasons: 1) if all samples for a given transcripts have 0 transcript counts, this transcript’s baseMean will be 0 and its LFC, p value, and padj will be set to NA; 2) if one replicate of a transcript is an outlier with an extreme count (detected by Cook’s distance), this transcript’s p value and padj will be set to NA; or 3) if a transcript is found to have a low mean normalized count after automatic independent filtering, this transcript’s padj will be set to NA.

## Acknowledgements

This work was supported in part by funding from the Defense Advanced Research Project Agency (DARPA) Safe Genes Program Grant (HR0011-17-2-0047) and an NIH New Innovator Award (1DP2AI152071-01) awarded to O.S.A.

## Materials and Data Availability

All constructs generated for this study are available on Addgene (https://www.addgene.org/) (Table S1). All fly lines created and/or used in this study are available at the Bloomington Drosophila Stock Center (https://bdsc.indiana.edu/) (Table S1). Raw sequencing data has been made publicly available at the NCBI SRA submission ID SUB6818910, BioProject PRJNA600654 (reviewer link https://dataview.ncbi.nlm.nih.gov/object/PRJNA600654?reviewer=uqk7ahvhcb93vp3jk7vkh3qiup).

## Author Contributions

O.S.A. conceived and designed the experiments. A.B. D.J.B, R.S., and T.Y. performed molecular and genetic experiments. P.H provided reagents and contributed to the experimental design. All authors contributed to the writing, analyzed the data, and approved the final manuscript.

## Competing Interests

O.S.A and A.B. have a patent pending on this technology. All other authors declare no competing interests.

## Supplemental Tables and Figures

**Table S1. Transgenic fly lines used in this study.** List of transgenic fly lines used in this study identifying the corresponding Addgene vector number, the Bloomington Drosophila Stock Center stock number, and the components of each integrated construct.

**Table S2. Complete data set for the Ubiq-CasRx and Ubiq-dCasRx bidirectional crosses.** Absolute counts of inheritance and phenotype penetrance for maternal and paternal inheritance of Ubiq-CasRx and Ubiq-dCasRx crosses to gRNA^array^-expressing flies. Each cross (paternal and maternal) was done in triplicate.

**Table S3. Complete data set for the Gal4/UASt-CasRx or Gal4/UASt-dCasRx crosses.** Absolute counts of inheritance and phenotype penetrance for the F_2_ generation resulting from F_1_ transheterozygote males expressing either UASt-CasRx/gRNA^array^ or UASt-dCasRx/gRNA^array^ crossed to Gal4 driver lines.

**Table S4. Illumina RNA sequencing whole-transcriptome analysis samples.** List of samples, in triplicate, analyzed for quantification of CasRx-mediated transcript reduction in comparison to dCasRx. The genotype, development stage or tissue type, and corresponding vectors are provided (Experimental = Ubiq-CasRx, Control = Ubiq-dCasRx).

**Table S5. Raw whole-transcriptome RNA-sequencing expression data.** Raw RNAseq data of all replicates analyzed in this study, identifying the expression of each gene in transcripts per million (TPM).

**Table S6. Raw whole-transcriptome RNA-sequencing count data.** Raw RNAseq data of all replicates analyzed in this study, quantifying the transcript counts for each gene.

**Table S7. RNA-sequencing expression data for all samples analyzed.** Expression data (TPM) for all analyzed genes and gRNAs across all DESeq2 data sets. The corresponding target gene of each sample (blue highlight) was used to analyze CasRx-mediated transcript reduction. The gRNA expression of the respective gene target is highlighted in green.

**Table S8. *GTP*-targeting DESeq2 analysis.** DESeq2 analysis of whole-transcriptome data for all three replicates comparing Ubiq-CasRx vs Ubiq-dCasRx for *GFP* targeting in first instar larvae.

**Table S9. *Notch-targeting* DESeq2 analysis.** DESeq2 analysis of whole-transcriptome data for all three replicates comparing Ubiq-CasRx vs Ubiq-dCasRx for *Notch* targeting in first instar larvae.

**Table S10. *yellow-targeting* DESeq2 analysis.** DESeq2 analysis of whole-transcriptome data for all three replicates comparing Ubiq-CasRx vs Ubiq-dCasRx for *yellow* targeting in embryos aged 17–20 hr.

**Table S11. *white-targeting* DESeq2 analysis.** DESeq2 analysis of whole-transcriptome data for all three replicates comparing Ubiq-CasRx vs Ubiq-dCasRx for *white* targeting in adult heads.

**Table S12. A list of CasRx-induced misexpressed transcripts based on the single-target DESeq2 analysis (Table S8–S11)**. The transcripts are annotated based on the FlyBase version FB2019_06. Transcripts affected by the presence of CasRx (padj < 0.05), also called the misexpressed transcripts, are labeled as 1, while transcripts not affected by the presence of CasRx (padj > 0.05) are labeled as 0. For the NA values, please refer to the Bioinformatics section in Methods.

**Table S13. Primers used for vector construction.** A list of primers and their respective sequences used to generate the constructs used in this study.

**Fig. S1.**
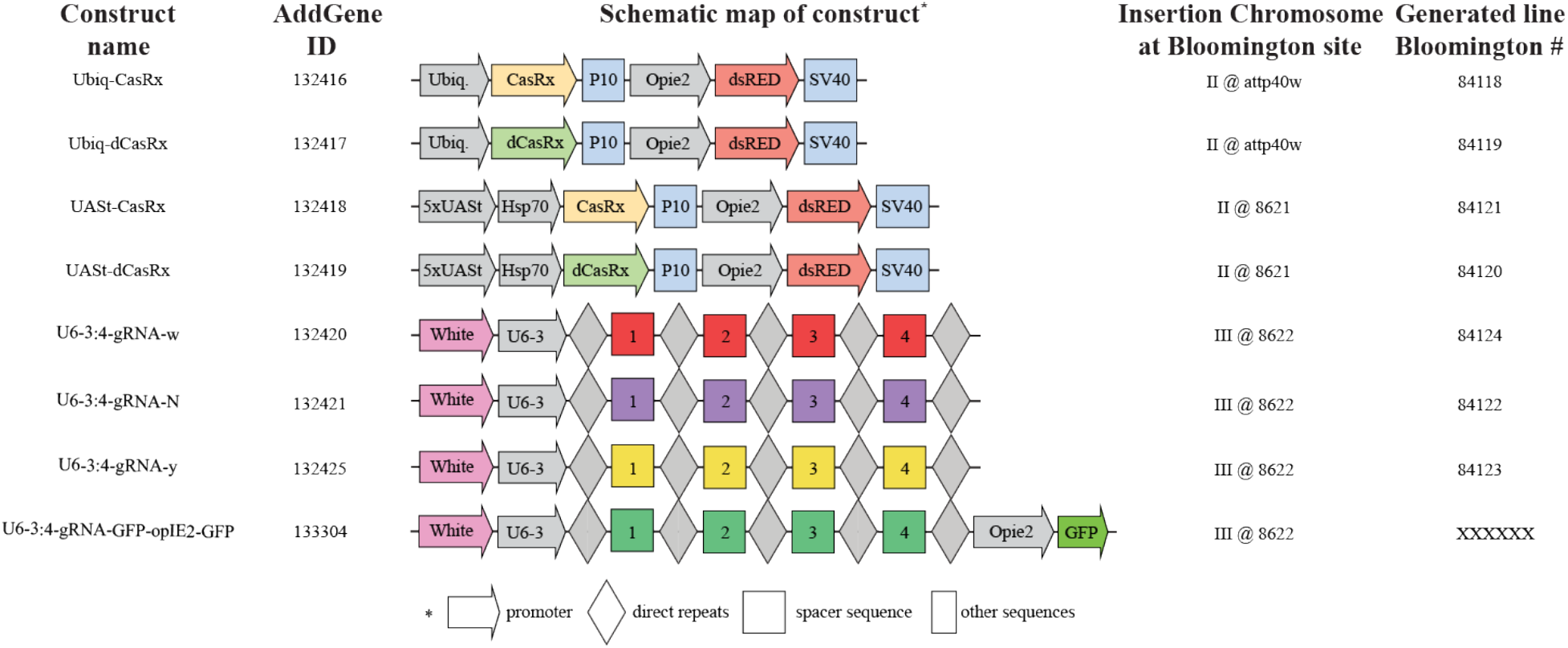
Schematic representation of constructs generated for this study. All constructs used in this study are depicted here along with addgene ID, insertion site, and Bloomington stock number.

**Fig. S2.**
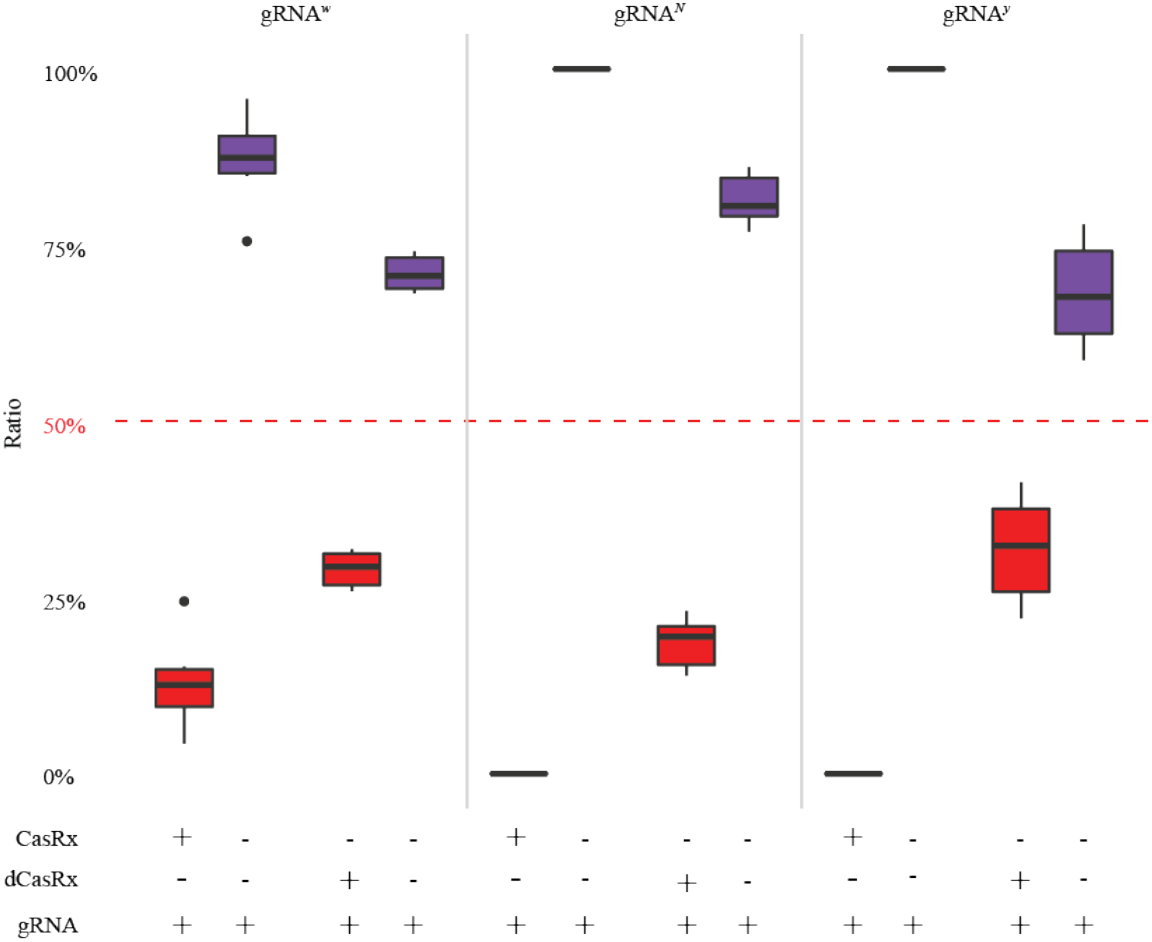
Complete inheritance data for bidirectional crosses. Complete inheritance plot for the bidirectional crosses featured in Fig 1. The plot includes all genotypes scored in all crosses between either Ubiq-CasRx or Ubiq-dCasRx and a respective gRNA^array^. In all crosses. gRNA^array^-only inheritance is dramatically higher than transheterozygote inheritance rates. including for Ubiq-dCasRx crosses.

**Fig. S3.**
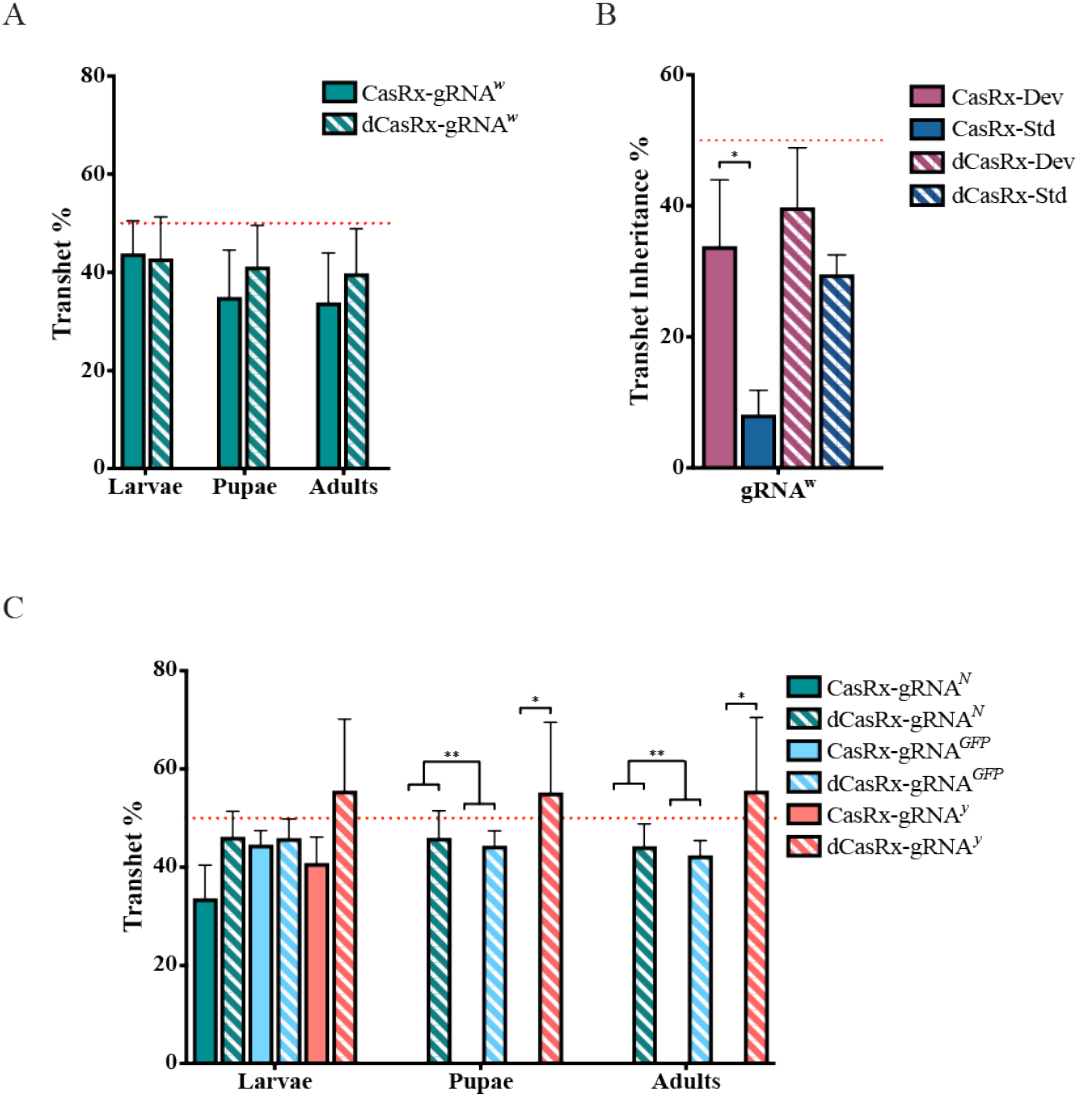
Development-related inheritance and lethality of Ubiq-CasRx and Ubiq-dCasRx transheterozygotes. (A-B) Transheterozygote percentages at larval, pupal, and adult developmental periods for each gRNA^array^ that produced an observable phenotype (*w*). There were no significant differences in inheritance. (C) Transheterozygote percentages through larval, pupal, and adult development periods for each gRNA^array^ that produced a lethal phenotype (*N, y, GFP*). No Ubiq-CasRx transheterozygotes developed beyond larvae.

**Fig. S4.**
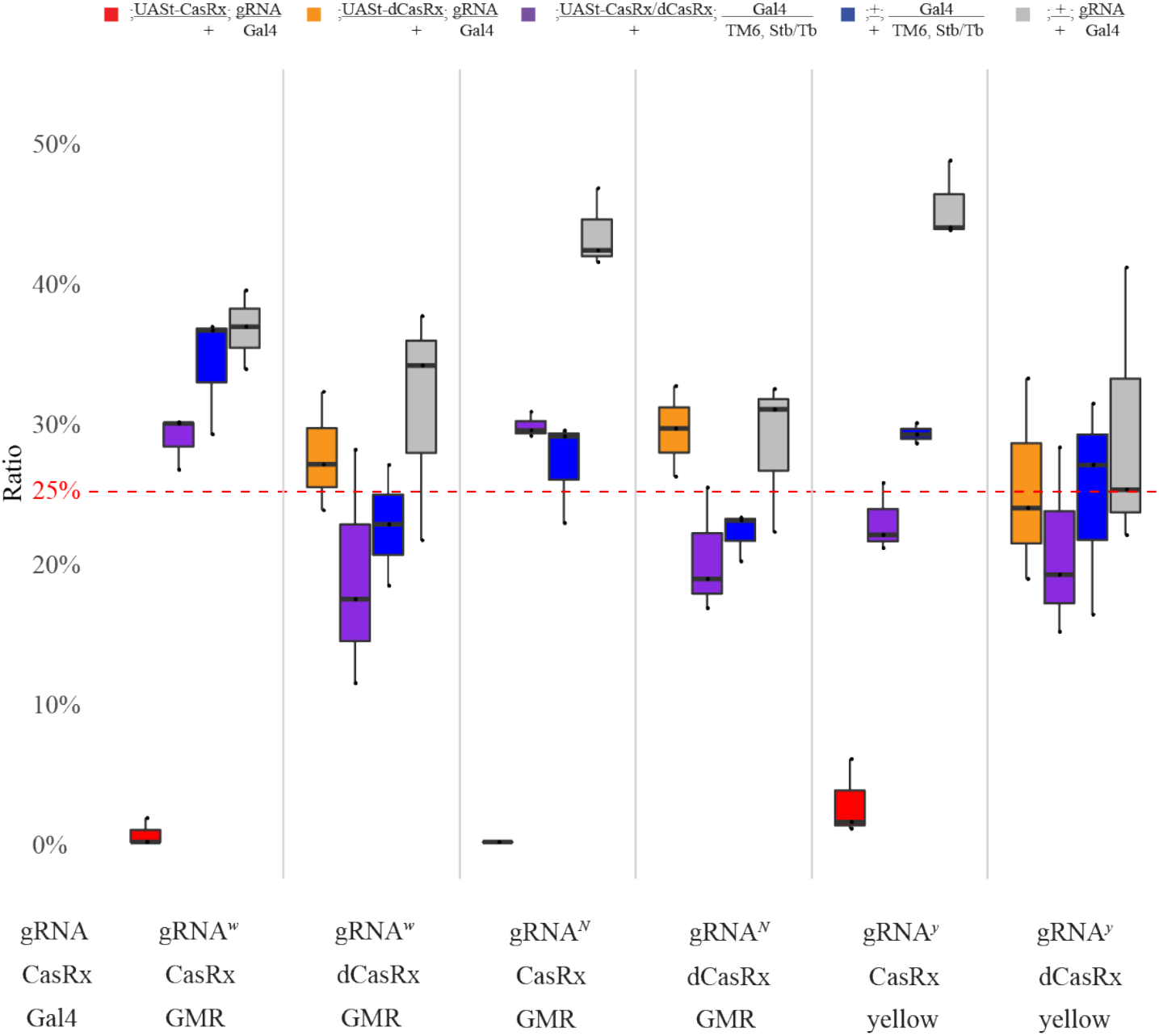
Complete inheritance data for binary Gal4/UAS crosses. The plot includes all genotypes scored in all crosses for UASt-CasRx and UASt-dCasRx. For all five gRNA^array^ targets, the inheritance of transheterozygous progeny expressing UASt-CasRx, a Gal4 driver, and a gRNA^array^ are lower compared to the other non-transheterozygous flies and to their corresponding dCasRx control group expressing UASt-dCasRx, a Gal4 driver, and a gRNA^array^.

**Fig. S5.**
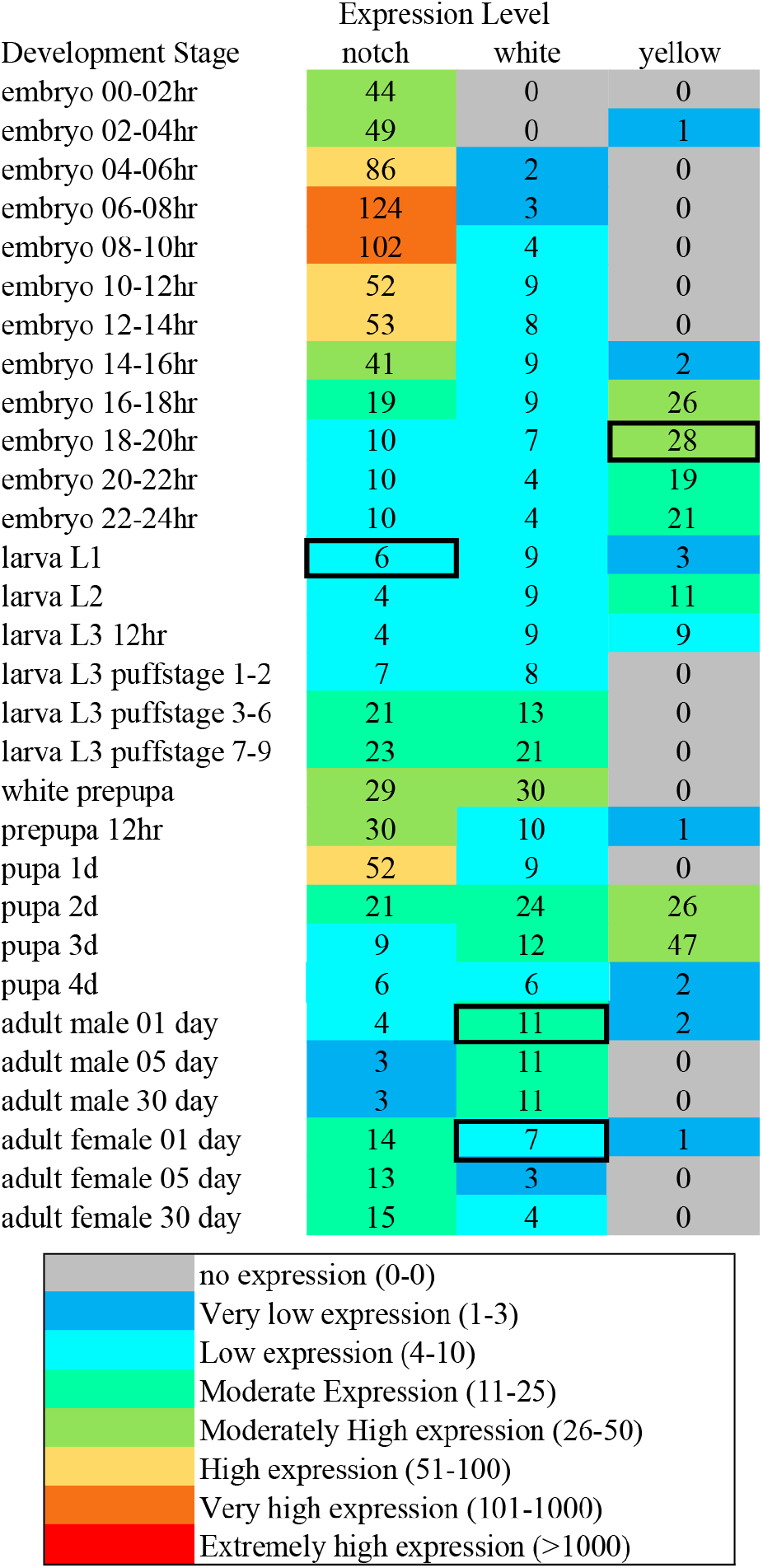
modENCODE transcript expression relative to *Drosophila melanogaster* development. Black box indicates which developmental period was chosen for RNA sequencing of samples for the analysis of CasRx-mediated transcript reduction in Ubiq-CasRx vs Ubiq-dCasRx comparison. Not included: GFP 1^st^ instar larvae were chosen for analysis of *GFP* transcript reduction.

**Fig. S6.**
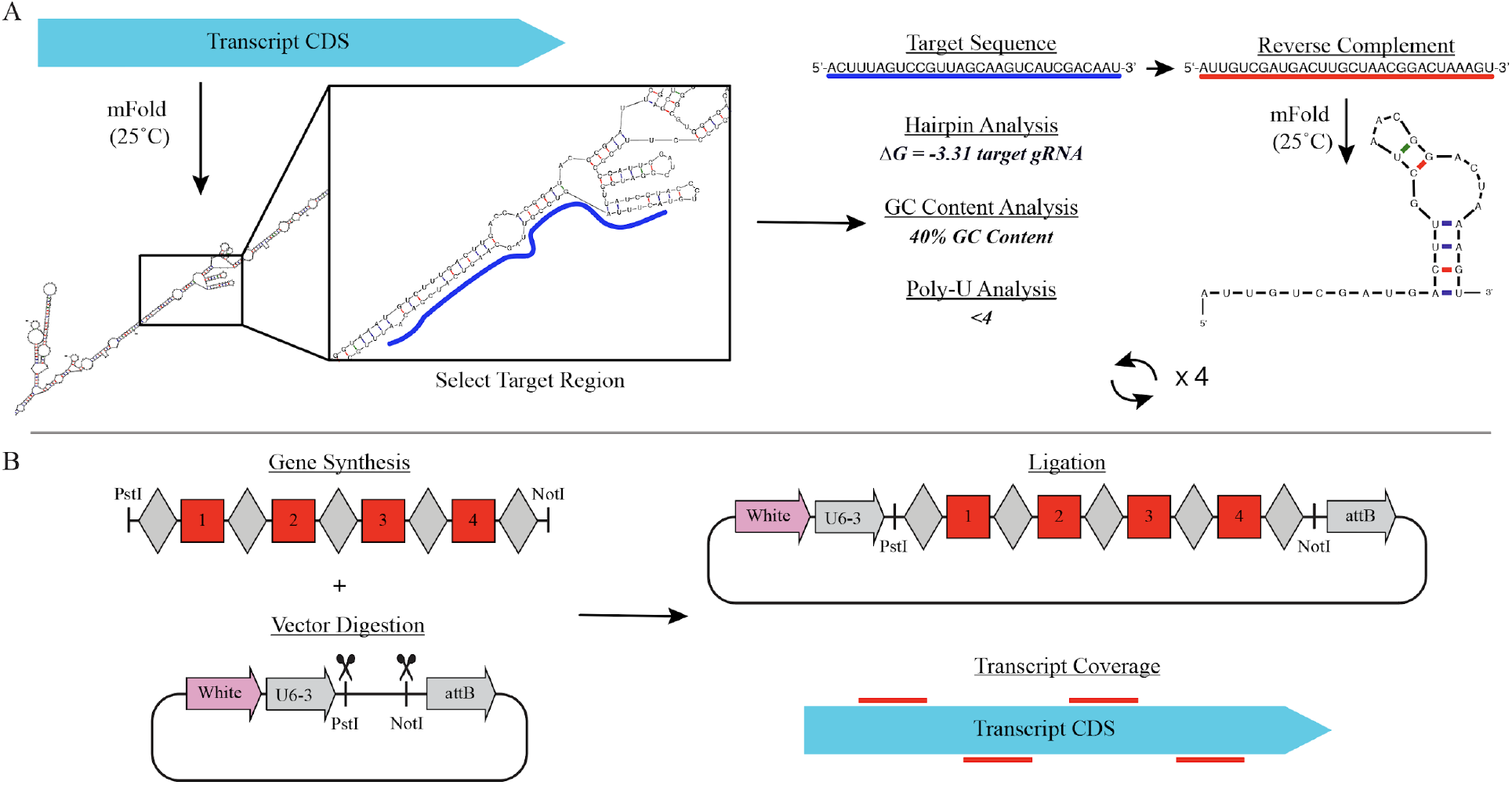
CasRx-gRNA^array^ transcript target selection and construct generation. (A) Schematic representing the workflow for gRNA choice. The transcript coding sequence (CDS) for a gene of interest (GOI) was entered into the mFold database (condition: 25°C) where predictive analysis identified the most probable secondary and tertiary folding of the entire transcript. We then chose specific regions that were predicted to be easily accessible for CasRx targeting (blue line), contained GC content between 30% and 70%, and possessed no poly-U stretches longer than 4 nt. We then converted the target sequence into the reverse complement (red line) and entered this spacer sequence into mFold (condition: 25°C) for hairpin analysis. This was repeated until four optimal target sites were selected. (B) Generation of gRNA^array^ construct. dsDNA was first synthesized to contain four spacer and five DR sequences with specific restriction sites present on the 5’ and 3’ end of the DNA. Simultaneously, the vector backbone containing the miniwhite marker, a U6:3 promoter fragment, and an attB site was digested using the corresponding restriction sites of the dsDNA gene fragment. The two pieces were then ligated together to generate a CasRx gRNA^array^ covering the majority of the transcript for the GOI.

## References

1. Adli M. The CRISPR tool kit for genome editing and beyond. Nat Commun. 2018;9: 1911.

2. Abudayyeh OO, Gootenberg JS, Konermann S, Joung J, Slaymaker IM, Cox DBT, et al. C2c2 is a single-component programmable RNA-guided RNA-targeting CRISPR effector. Science. 2016;353: aaf5573.

3. East-Seletsky A, O’Connell MR, Burstein D, Knott GJ, Doudna JA. RNA Targeting by Functionally Orthogonal Type VI-A CRISPR-Cas Enzymes. Molecular Cell. 2017. pp. 373–383.e3. doi:10.1016/j.molcel.2017.04.008

4. Konermann S, Lotfy P, Brideau NJ, Oki J, Shokhirev MN, Hsu PD. Transcriptome Engineering with RNA-Targeting Type VI-D CRISPR Effectors. Cell. 2018;173: 665–676.e14.

5. Kushawah G, del Prado JA-N, Martinez-Morales JR, DeVore M, Guelfo JR, Brannan EO, et al. CRISPR-Cas13d induces efficient mRNA knock-down in animal embryos. doi:10.1101/2020.01.13.904763

6. Abudayyeh OO, Gootenberg JS, Essletzbichler P, Han S, Joung J, Belanto JJ, et al. RNA targeting with CRISPR-Cas13. Nature. 2017;550: 280–284.

7. Smargon AA, Cox DBT, Pyzocha NK, Zheng K, Slaymaker IM, Gootenberg JS, et al. Cas13b Is a Type VI-B CRISPR-Associated RNA-Guided RNase Differentially Regulated by Accessory Proteins Csx27 and Csx28. Mol Cell. 2017;65: 618–630.e7.

8. East-Seletsky A, O’Connell MR, Knight SC, Burstein D, Cate JHD, Tjian R, et al. Two distinct RNase activities of CRISPR-C2c2 enable guide-RNA processing and RNA detection. Nature. 2016;538: 270–273.

9. Yan WX, Chong S, Zhang H, Makarova KS, Koonin EV, Cheng DR, et al. Cas13d Is a Compact RNA-Targeting Type VI CRISPR Effector Positively Modulated by a WYL-Domain-Containing Accessory Protein. Mol Cell. 2018;70: 327–339.e5.

10. Perrimon N, Ni J-Q, Perkins L. In vivo RNAi: today and tomorrow. Cold Spring Harb Perspect Biol. 2010;2: a003640.

11. Champer J, Buchman A, Akbari OS. Cheating evolution: engineering gene drives to manipulate the fate of wild populations. Nat Rev Genet. 2016;17: 146–159.

12. Buchman A, Gamez S, Li M, Antoshechkin I, Li H-H, Wang H-W, et al. Broad dengue neutralization in mosquitoes expressing an engineered antibody. PLoS Pathog. 2020;16: e1008103.

13. Mathur G, Sanchez-Vargas I, Alvarez D, Olson KE, Marinotti O, James AA. Transgene-mediated suppression of dengue viruses in the salivary glands of the yellow fever mosquito, Aedes aegypti. Insect Mol Biol. 2010;19: 753–763.

14. Franz AWE, Sanchez-Vargas I, Adelman ZN, Blair CD, Beaty BJ, James AA, et al. Engineering RNA interference-based resistance to dengue virus type 2 in genetically modified Aedes aegypti. Proc Natl Acad Sci U S A. 2006;103: 4198–4203.

15. Yen P-S, James A, Li J-C, Chen C-H, Failloux A-B. Synthetic miRNAs induce dual arboviral-resistance phenotypes in the vector mosquito Aedes aegypti. Communications Biology. 2018;1: 11.

16. Buchman A, Gamez S, Li M, Antoshechkin I, Li H-H, Wang H-W, et al. Engineered resistance to Zika virus in transgenic Aedes aegypti expressing a polycistronic cluster of synthetic small RNAs. Proc Natl Acad Sci U S A. 2019. doi:10.1073/pnas.1810771116

17. Dietzl G, Chen D, Schnorrer F, Su K-C, Barinova Y, Fellner M, et al. A genome-wide transgenic RNAi library for conditional gene inactivation in Drosophila. Nature. 2007;448: 151–156.

18. Ni J-Q, Liu L-P, Binari R, Hardy R, Shim H-S, Cavallaro A, et al. A Drosophila resource of transgenic RNAi lines for neurogenetics. Genetics. 2009;182: 1089–1100.

19. Ni J-Q, Zhou R, Czech B, Liu L-P, Holderbaum L, Yang-Zhou D, et al. A genome-scale shRNA resource for transgenic RNAi in Drosophila. Nat Methods. 2011;8: 405–407.

20. Ni J-Q, Markstein M, Binari R, Pfeiffer B, Liu L-P, Villalta C, et al. Vector and parameters for targeted transgenic RNA interference in Drosophila melanogaster. Nat Methods. 2008;5: 49–51.

21. Heigwer F, Port F, Boutros M. RNA Interference (RNAi) Screening in Drosophila. Genetics. 2018;208: 853–874.

22. Kulkarni MM, Booker M, Silver SJ, Friedman A, Hong P, Perrimon N, et al. Evidence of off-target effects associated with long dsRNAs in Drosophila melanogaster cell-based assays. Nature Methods. 2006. pp. 833–838. doi:10.1038/nmeth935

23. Ma Y, Creanga A, Lum L, Beachy PA. Prevalence of off-target effects in Drosophila RNA interference screens. Nature. 2006;443: 359–363.

24. Perrimon N, Mathey-Prevot B. Matter arising: off-targets and genome-scale RNAi screens in Drosophila. Fly. 2007;1: 1–5.

25. Markstein M, Pitsouli C, Villalta C, Celniker SE, Perrimon N. Exploiting position effects and the gypsy retrovirus insulator to engineer precisely expressed transgenes. Nat Genet. 2008;40: 476–483.

26. Chakraborty C, Teoh SL, Das S. The Smart Programmable CRISPR Technology: A Next Generation Genome Editing Tool for Investigators. Curr Drug Targets. 2017;18: 1653–1663.

27. Jinek M, Chylinski K, Fonfara I, Hauer M, Doudna JA, Charpentier E. A programmable dual-RNA-guided DNA endonuclease in adaptive bacterial immunity. Science. 2012;337: 816–821.

28. Akbari OS, Oliver D, Eyer K, Pai C-Y. An Entry/Gateway cloning system for general expression of genes with molecular tags in Drosophila melanogaster. BMC Cell Biol. 2009;10: 8.

29. Biessmann H. Molecular analysis of the yellow gene (y) region of Drosophila melanogaster. Proceedings of the National Academy of Sciences. 1985. pp. 7369–7373. doi:10.1073/pnas.82.21.7369

30. Sullivan DT, Sullivan MC. Transport defects as the physiological basis for eye color mutants of Drosophila melanogaster. Biochemical Genetics. 1975. pp. 603–613. doi:10.1007/bf00484918

31. Kidd S, Kelley MR, Young MW. Sequence of the notch locus of Drosophila melanogaster: relationship of the encoded protein to mammalian clotting and growth factors. Molecular and Cellular Biology. 1986. pp. 3094–3108. doi:10.1128/mcb.6.9.3094

32. Lindsley DL. Genetic Variations of Drosophila Melanogaster [by] Dan L. Lindsley and E.H. Grell. 1968.

33. Kandul NP, Liu J, Sanchez CHM, Wu SL, Marshall JM, Akbari OS. Transforming insect population control with precision guided sterile males with demonstration in flies. Nat Commun. 2019;10: 84.

34. Port F, Chen H-M, Lee T, Bullock SL. Optimized CRISPR/Cas tools for efficient germline and somatic genome engineering in Drosophila. Proc Natl Acad Sci U S A. 2014;111: E2967–76.

35. Micchelli CA, Perrimon N. Evidence that stem cells reside in the adult Drosophila midgut epithelium. Nature. 2006;439: 475–479.

36. Leonardi J, Fernandez-Valdivia R, Li Y-D, Simcox AA, Jafar-Nejad H. Multiple O-glucosylation sites on Notch function as a buffer against temperature-dependent loss of signaling. Development. 2011;138: 3569–3578.

37. Simón R, Aparicio R, Housden BE, Bray S, Busturia A. Drosophila p53 controls Notch expression and balances apoptosis and proliferation. Apoptosis. 2014;19: 1430–1443.

38. Geyer PK, Green MM, Corces VG. Tissue-specific transcriptional enhancers may act in trans on the gene located in the homologous chromosome: the molecular basis of transvection in Drosophila. EMBO J. 1990;9: 2247–2256.

39. Brand AH, Perrimon N. Targeted gene expression as a means of altering cell fates and generating dominant phenotypes. Development. 1993;118: 401–415.

40. Saj A, Arziman Z, Stempfle D, van Belle W, Sauder U, Horn T, et al. A combined ex vivo and in vivo RNAi screen for notch regulators in Drosophila reveals an extensive notch interaction network. Dev Cell. 2010;18: 862–876.

41. Massey JH, Chung D, Siwanowicz I, Stern DL, Wittkopp PJ. The yellow gene influences Drosophila male mating success through sex comb melanization. bioRxiv. 2019. p. 673756. doi:10.1101/673756

42. Pfeifer TA, Hegedus DD, Grigliatti TA, Theilmann DA. Baculovirus immediate-early promoter-mediated expression of the Zeocin™ resistance gene for use as a dominant selectable marker in Dipteran and Lepidopteran insect cell lines. Gene. 1997;188: 183–190.

43. Graveley BR, Brooks AN, Carlson JW, Duff MO, Landolin JM, Yang L, et al. The developmental transcriptome of Drosophila melanogaster. Nature. 2011;471: 473–479.

44. Love MI, Huber W, Anders S. Moderated estimation of fold change and dispersion for RNA-seq data with DESeq2. Genome Biol. 2014;15: 550.

45. Meeske AJ, Nakandakari-Higa S, Marraffini LA. Cas13-induced cellular dormancy prevents the rise of CRISPR-resistant bacteriophage. Nature. 2019;570: 241–245.

46. Peretz G, Bakhrat A, Abdu U. Expression of the Drosophila melanogaster GADD45 homolog (CG11086) affects egg asymmetric development that is mediated by the c-Jun N-terminal kinase pathway. Genetics. 2007;177: 1691–1702.

47. Wessels H-H, Méndez-Mancilla A, Guo X, Legut M, Daniloski Z, Sanjana NE. Principles for rational Cas13d guide design. doi:10.1101/2019.12.27.889089

48. Zuker M. Mfold web server for nucleic acid folding and hybridization prediction. Nucleic Acids Res. 2003;31: 3406–3415.

49. Gibson DG, Young L, Chuang R-Y, Venter JC, Hutchison CA 3rd, Smith HO. Enzymatic assembly of DNA molecules up to several hundred kilobases. Nat Methods. 2009;6: 343–345.

50. Pfeiffer BD, Truman JW, Rubin GM. Using translational enhancers to increase transgene expression in Drosophila. Proc Natl Acad Sci U S A. 2012;109: 6626–6631.

51. Gamez S, Antoshechkin I, Mendez-Sanchez SC, Akbari OS. The Developmental Transcriptome of Ae. albopictus, a Major Worldwide Human Disease Vector. bioRxiv. 2019. p. 753962. doi:10.1101/753962

52. Dobin A, Davis CA, Schlesinger F, Drenkow J, Zaleski C, Jha S, et al. STAR: ultrafast universal RNA-seq aligner. Bioinformatics. 2013. pp. 15–21. doi:10.1093/bioinformatics/bts635

53. Liao Y, Smyth GK, Shi W. featureCounts: an efficient general purpose program for assigning sequence reads to genomic features. Bioinformatics. 2014. pp. 923–930. doi:10.1093/bioinformatics/btt656

